# Blue light irradiation induces pollen tube rupture in various flowering plants

**DOI:** 10.1101/2023.12.06.570347

**Authors:** Naoya Sugi, Daichi Susaki, Yoko Mizuta, Tetsu Kinoshita, Daisuke Maruyama

**Affiliations:** Kihara Institute for Biological Research, Yokohama City University, Maioka-cho, Totsuka-ku, Yokohama, Kanagawa, 244-0813, Japan; Institute for Advanced Research (IAR), Nagoya University, Furo-cho, Chikusa-ku, Nagoya, Aichi 464-8601, Japan; Institute of Transformative Bio-Molecules (WPI-ITbM), Nagoya University, Furo-cho, Chikusa-ku, Nagoya, Aichi 464-8601, Japan

**Author notes:** **Corresponding Author:** N. Sugi, Kihara Institute for Biological Research, Yokohama City University, Maioka-cho, Totsuka-ku, Yokohama, Kanagawa, 244-0813, Japan., (E-mail).

**Keywords:** blue light irradiation, pollen tube rupture, apical growth, calcium, cell wall integrity, Arabidopsis

## Abstract

Pollen tubes exhibit one of the fastest apical growth rates among plant cells. Maintaining the proper balance between turgor pressure and cell wall synthesis at the pollen tube tip is crucial for this rapid growth, and any disruption can result in pollen tube rupture. In our study, we reveal that exposure to short-wavelength visible light, specifically blue light, induces pollen tube rupture. The frequency of pollen tube rupture increases in an intensity-dependent manner. Additionally, we observed Ca^2+^ influx after blue light irradiation, accompanying with either pollen tube rupture or a temporary halt in elongation. These findings offer insights into the interplay between pollen tube integrity maintenance and Ca^2+^ influx at the pollen tube tip, presenting a novel and efficient method to control pollen tube burst.

**Subject Areas:** (1) growth and development

(11) new methodology

## Introduction

In flowering plants, the pollen tube serves as a vehicle to transport two nonmotile sperm cells. It is the fastest-growing plant cell, elongating at a rate exceeding 5 μm/min in *Arabidopsis* (Cheung et al., 2010). Upon contact with the stigma, pollen grains hydrate and initiate pollen tube germination. The pollen tube extends through female tissues, ultimately reaching one of two synergid cells in an ovule. At this point, the tube ruptures, releasing two sperm cells. These steps are essential to accomplish double fertilization.

During pollen tube growth, maintaining the integrity of the pollen tube cell wall is crucial and is tightly regulated by various factors (Ogawa and Kessler, 2023). Catharanthus roseus receptor-like kinase 1-like (CrRLK1L)-type receptors ANXUR1/2 (ANX1/2) and BUDDHA PAPER SEAL1/2 (BUPS1/2), along with GPI-anchored proteins LORELEI-LIKE GENE2/3 (LLG2/3), localize to the plasma membrane at the pollen tube tip. This receptor complex interacts with ligands RAPID ALKALINIZATION FACTOR 4/19 (RALF4/19), secreted from the pollen tube tip. This autocrine signaling stabilizes pollen tube integrity. Ca^2+^ channels MILDEW RESISTANCE LOCUS O 1/5/9/15 (MLO1/5/9/15), downstream of the ANX/BUPS/LLG/RALF pathway, play a crucial role, and mutants in this pathway exhibit spontaneous pollen tube rupture due to the failure of Ca^2+^ gradient formation at the tube tip (Gao et al., 2023). LEUCINE-RICH REPEAT EXTENSINs (LRXs) also interact with RALF peptides. This LRX-RALF interaction contributes to cell wall integrity through a condensing effect. Pollen tubes lacking RALF4/19 or LRXs display abnormalities in cell wall structure, resulting in bursting or growth defects (Fabrice et al., 2018, Moussu et al., 2023). Considering the physicochemical state of pollen tube growth and the maintenance of cell wall integrity, turgor pressure serves as a vital driving force for apical growth, and the viscous liquid film created by cell wall materials at the pollen tube tip is an extremely unstable state. Consequently, pollen tube burst is believed to be ultimately driven by an imbalance between cell wall strength and cytosolic turgor pressure (Hill et al., 2012, Weigand et al., 2023). In summary, the cooperatively controlled regulation of pollen tube cell wall through the autocrine ANX/BUPS/LLG/RALF signaling pathway is essential for maintaining pollen tube integrity (Dehors et al., 2019, Hill et al., 2012, McKenna et al., 2009, Kroeger et al., 2011, Ogawa and Kessler, 2023).

After pollen tube burst, double fertilization is completed within 10 min (Hamamura et al., 2011). Swift removal of the inner vegetative plasma membrane (IVPM) and relocation of GENERATIVE CELL SPECIFIC 1 (GCS1)/HAPLESS 2 (HAP2) must be achieved before gamete fusion following pollen tube burst (Sugi et al., 2023, Sprunck et al., 2012, Wang et al., 2022). Analyzing these membrane dynamics poses a significant challenge due to the need for observations in seconds or milliseconds. In the current scenario, inducing pollen tube burst in *in vitro* conditions requires the treatment of a liquid medium containing recombinant RALF34 peptide, reactive oxygen species (ROS), or osmotic stress. Therefore, a moderate technique to control the timing of pollen tube burst has not yet been established. The technology to control the function of a specific cell using light irradiation is compatible with time-lapse imaging by a microscope and has brought about breakthroughs in cell biology and embryology. These include laser ablation (Susaki et al., 2015, Nagahara et al., 2020, Higashiyama et al., 2001), infrared laser-evoked gene operator (Kamei et al., 2009), optogenetic tools (Zhou et al., 2021, He et al., 2021), and fluorescence recovery after photobleaching (Swaminathan, 1997). Despite the risk of phototoxicity, these have contributed to the progress of life science. Here, we report that a very simple treatment, blue light irradiation to the pollen tube tips, induces pollen tube rupture. This new method may aid in understanding the precise control of pollen tube tip integrity and provide a novel option for research in the field of plant reproduction.

## Results

### Blue light irradiation induces pollen tube rupture

Through hundreds of fluorescent observations of growing *Arabidopsis* pollen tubes, we observed that pollen tubes tend to rupture during excitation light irradiation. To characterize this phenomenon, we investigated the relationship between the wavelength and duration of light irradiation and rupture using *Arabidopsis* pollen tubes. When pollen tubes of wild-type (WT) plants were continuously irradiated with excitation light of the CFP filter set (blue light in the 425–445 nm wavelength range), approximately 60% of the pollen tubes burst (Figure 1A, Supplementary Movie 1). We then observed the accumulated frequency of pollen tube ruptures over time. The earliest rupture was observed at 16 s after the start of irradiation, with most ruptures occurring within 1 min (Figure 1B). Frequent pollen tube rupture could be induced even with a restricted duration of light irradiation to 15 s (Figure 1C, Supplementary Movie 3). Furthermore, the frequency of pollen tube rupture increased with excitation light intensity in a dose-dependent manner (Figure 1C, Supplementary Movie 2). To investigate wavelength dependence, we examined the response to excitation light irradiation with GFP and RFP filter sets for 15 s. The results showed that excitation light from GFP and RFP filters did not induce pollen tube burst (Supplementary Movie 4,5), indicating that blue light can induce these phenomena.

**Figure 1.**
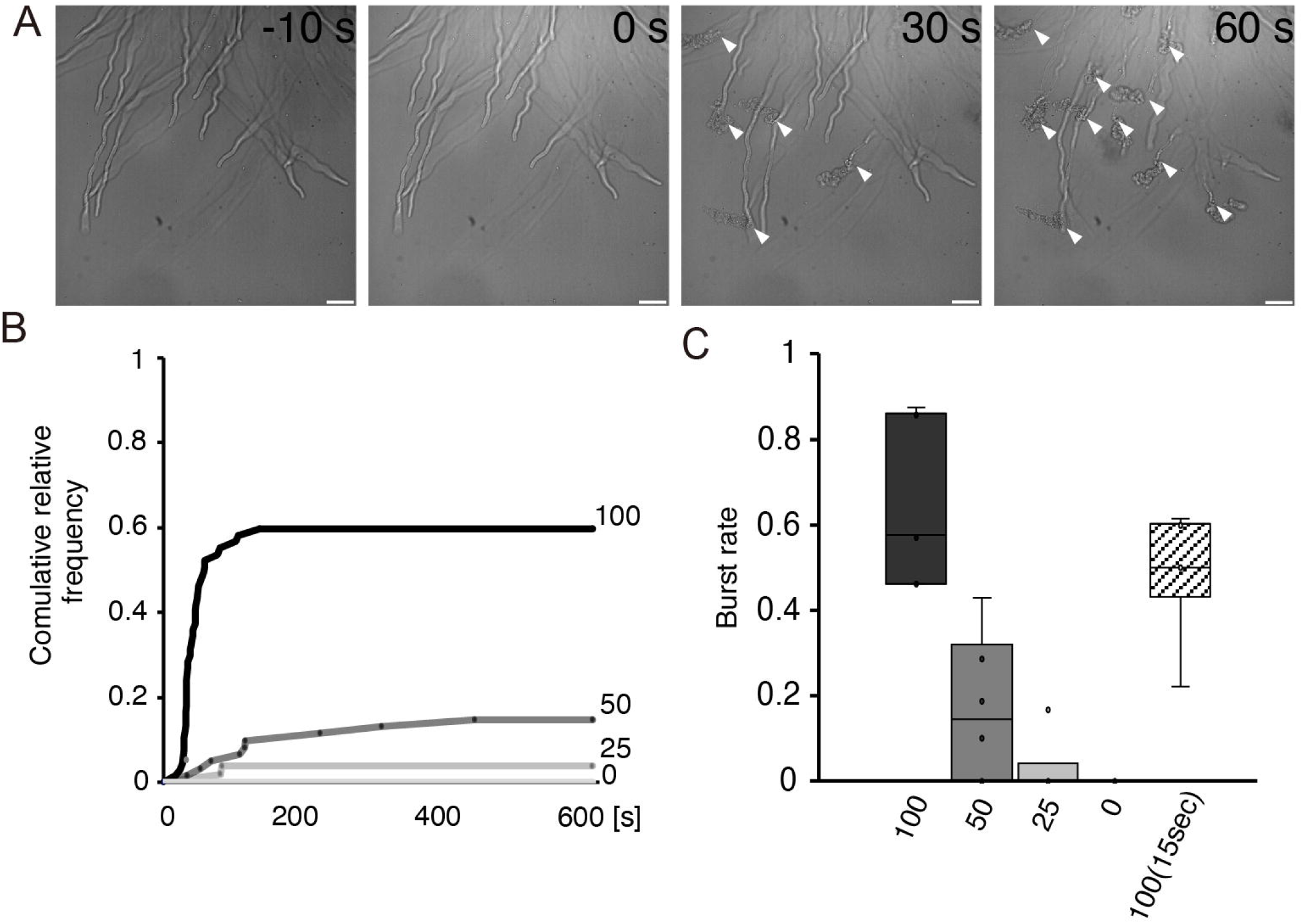
Blue light irradiation induced pollen tube burst. (A) Montaged images of pollen tubes before and after blue light irradiation. (B) Cumulative relative frequency of pollen tube burst in different light intensities. (C) Pollen tube burst rate in different light intensities in 5 min. Filled arrowheads indicate pollen tube burst. `Box plots represent the means with 25th and 75th percentiles with minimum and maximum whiskers.` Scale bars: 30 μm.

### Blue light irradiation could induce pollen tube rupture in Torenia and Tobacco

To determine whether blue light-induced rupture is a common feature of growing pollen tubes in various plant species, we challenged pollen tubes from *Nicotiana benthamiana* and *Torenia fournieri* with blue light irradiation. *N. benthamiana* produces bicellular pollen containing a single generative cell in the pollen vegetative cell, and the generative cell divides into two sperm cells during pollen tube growth. The width of the pollen tube in *N. benthamiana* is approximately 10 μm, twice as thick as those in *Arabidopsis*. When *N. benthamiana* pollen tubes were irradiated with blue light after emerging into the growth medium from the cut style, abrupt rupturing was observed within 2 min (54%, n = 13, Figure 2A). A similar blue light irradiation test was performed in *T. fournieri*, which produces bicellular pollen with an intermediate diameter of the pollen tube (∼8 μm). Pollen tube rupture was also observed in this organism (50%, n = 38, Figure 2B). These results suggest that blue light-induced pollen tube rupture is not limited to *Arabidopsis* and may be a common phenomenon across plant species despite of their morphological variations in pollen tubes.

**Figure 2.**
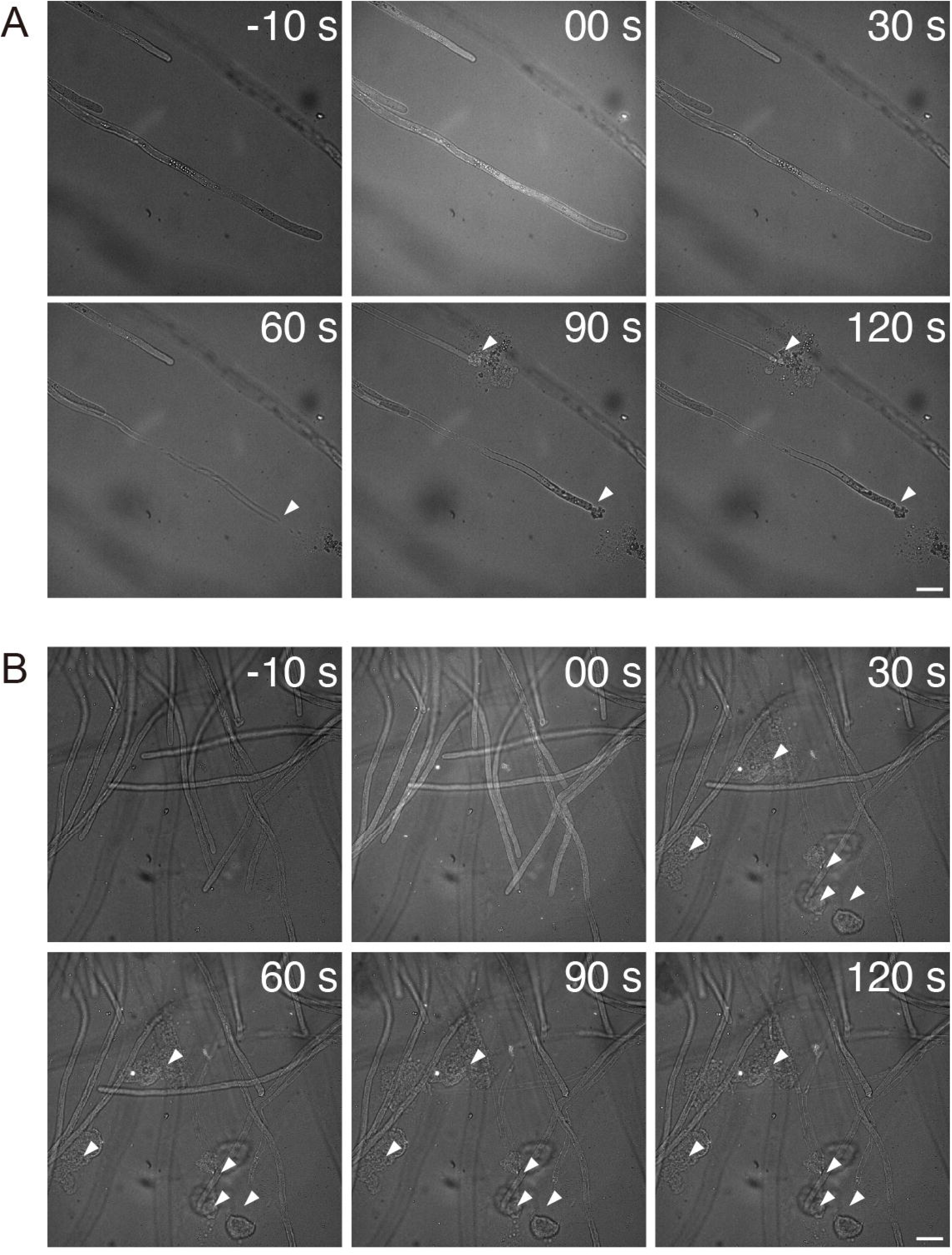
Blue light irradiation in Tobacco and Torenia. (A) Montaged images of pollen tubes in Tobacco. (B) Montaged images of pollen tubes in Torenia. Filled arrowheads indicate pollen tube burst. Scale bars: 30 μm.

### Blue light irradiation caused Ca^2+^ influx from pollen tube tip

In *Arabidopsis*, calcium dynamics play crucial roles in maintaining cell wall integrity during pollen tube growth (Duan et al., 2014; Gao et al., 2023). To monitor Ca^2+^ dynamics under blue light irradiation, we observed pollen tubes expressing the calcium sensor protein GCaMP (Nakai et al., 2001). Before blue light irradiation, we detected a calcium maximum at the subapical region, gradually decreasing toward the shank region (Figure 3A). Throughout the time-lapse imaging, the apical Ca^2+^ gradient was maintained, and pollen tubes elongated at a constant rate (Figure 3 D). To simultaneously observe the fluorescence pattern of GCaMP under blue light conditions, a microscope system was constructed to irradiate the excitation light of the CFP filter while observing confocal microscopy with a 488 nm laser. When this system was used to irradiate pollen tubes with blue light for 15 s, the high fluorescent intensity region extended to the basal side in about 90% of the pollen tubes, suggesting that blue light irradiation caused Ca^2+^ influx (Figure 3 BC, EF). Among them, 34% (n=171) exhibited pollen tube rupture during or after the Ca^2+^ influx. The remaining 66% (n=171) did not rupture and showed a significant inhibition of subsequent pollen tube elongation (Figure 3 F). Interestingly, the pollen tube displayed temporal loss and partial reformation of the Ca^2+^ gradient during the unusual growth arrest. This suggests that blue light irradiation may induce stochastic pollen tube rupture by acting on Ca^2+^-mediated regulation of cell wall flexibility at the pollen tube tip.

**Figure 3.**
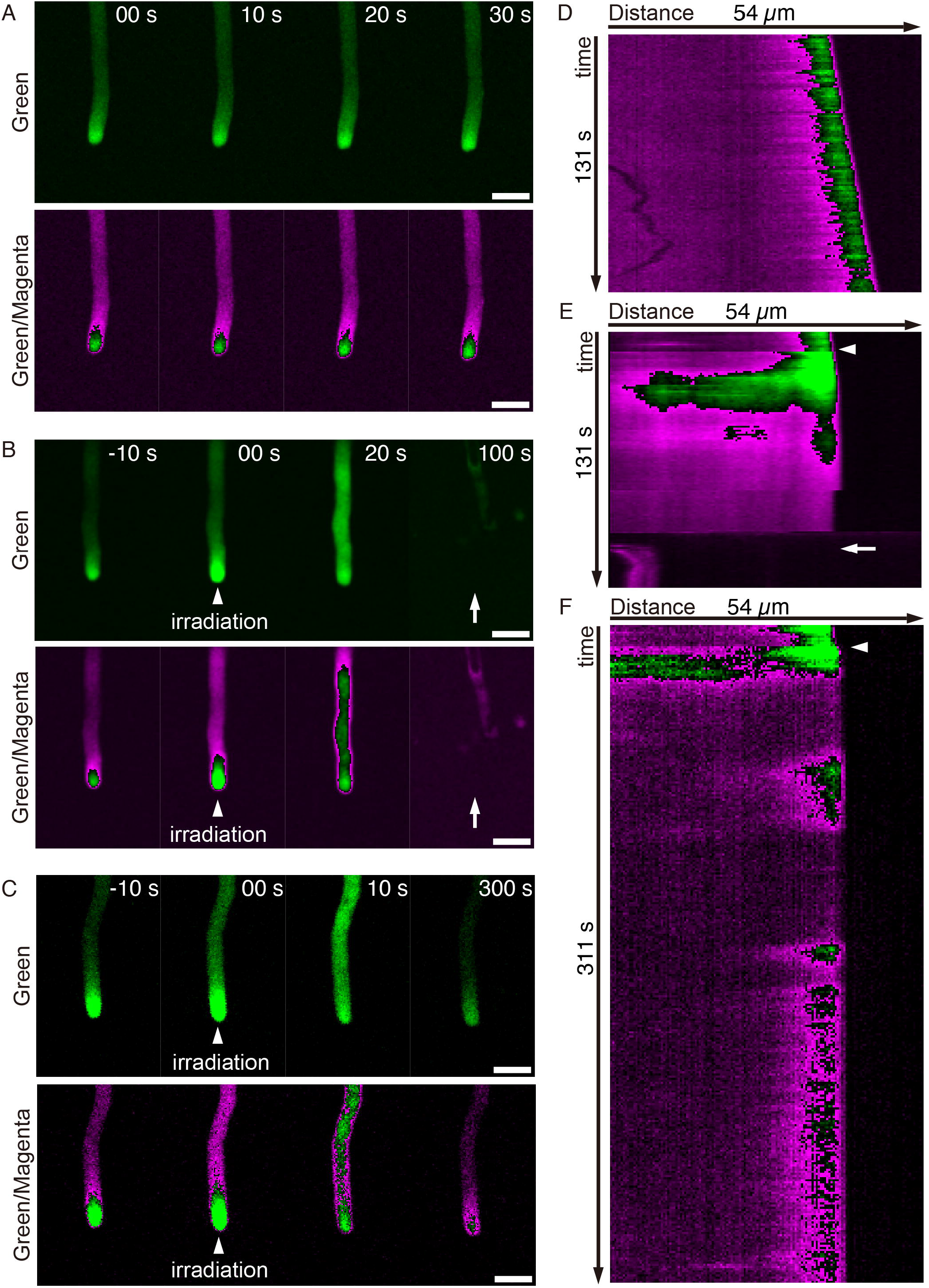
Calcium imaging during blue light irradiation. (A–C) Montaged images of pollen tubes expressing GCaMP; steady state in green channel (A, upper panel), steady state in green/magenta heatmap (A, lower panel), pollen tube burst after blue light irradiation in green channel (B, upper panel), pollen tube burst after blue light irradiation in green/magenta heatmap (B, lower panel), pollen tube growth arrest after blue light irradiation in green channel (C, upper panel), pollen tube growth arrest after blue light irradiation in green/magenta heatmap (C, lower panel). (D–F) Kymograph of pollen tubes expressing GCaMP in green/magenta heatmap; steady state (D), pollen tube burst after blue light irradiation (E), pollen tube growth arrest after blue light irradiation (F). Arrows indicate pollen tube burst; filled arrowheads show the starting time of blue light irradiation. Scale bars: 10 μm.

### Apical region was susceptible for blue light stimuli

Pollen tube rupture was induced using blue light irradiation covering the entire field of view, stimulating a wide range of the pollen tube: tip and other basal shank regions. To elucidate the pollen tube’s susceptibility to light stimuli, a partial blue light irradiation was done. The field of blue light irradiation was restricted by narrowing the aperture of the microscope (Figure 4 A–C, dotted circle). We observed a 23% (n = 85, Figure 4D) rupture in pollen tubes stimulated, including the tip region, comparable to the initial condition using wide-range irradiation (Figure 4C, open arrowheads; Figure 1A). However, 0% (n = 72, Figure 4D) of the pollen tubes ruptured when the blue light irradiated only the shank but not the tip region (Figure 4C, arrows). These observations indicate a distinct sensitivity of the pollen tube apical region to blue light irradiation.

**Figure 4.**
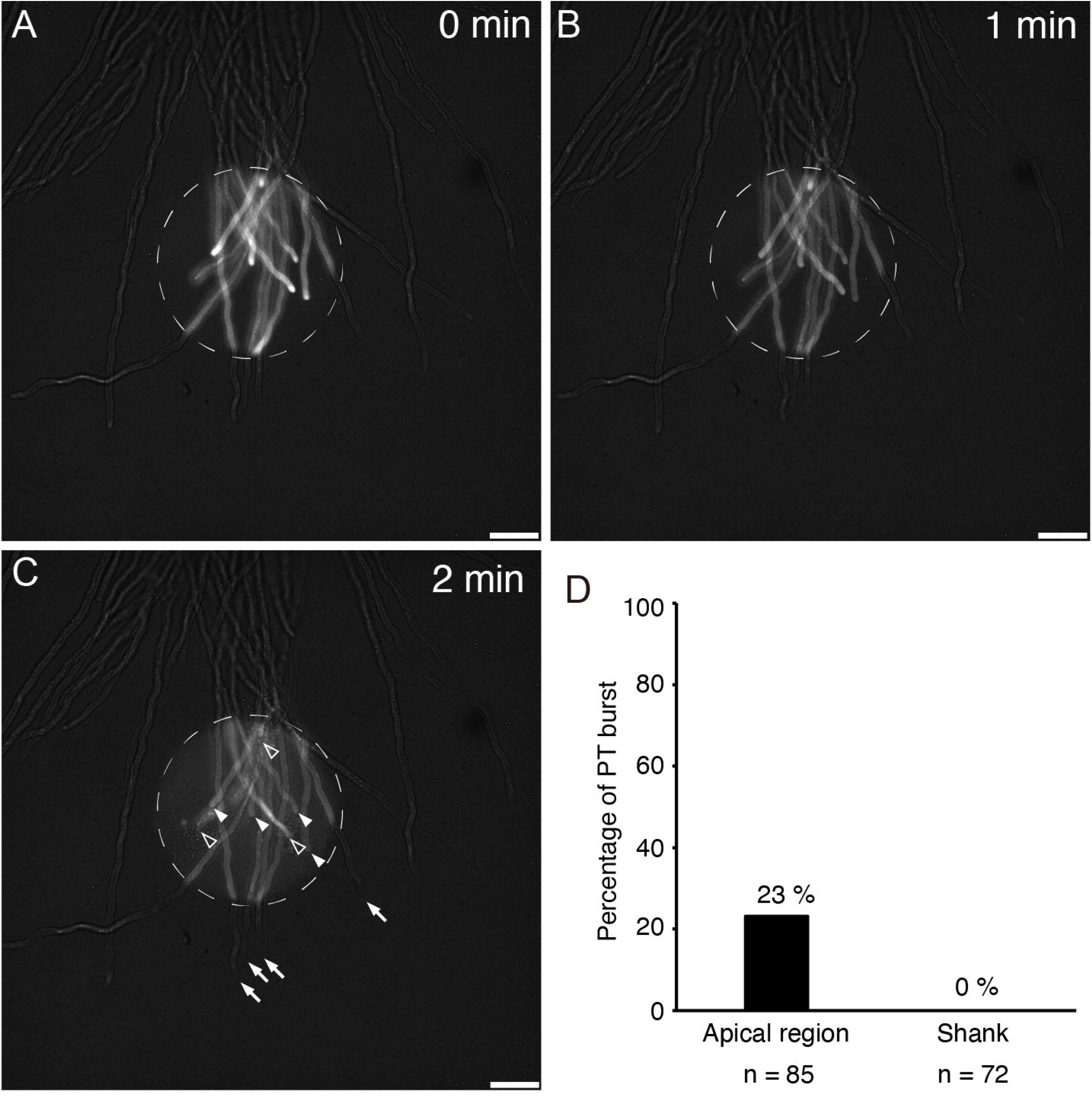
Limited irradiation of blue light to the pollen tubes. (A–C) Montaged images of pollen tubes expressing GCaMP with limited blue light irradiation. (D) Percentage of pollen tube burst in two pollen tube types; the pollen tube tip was included in irradiated area (Apical region), and only the pollen tube shank was included in irradiated area (shank). Arrows indicate pollen tube burst. Scale bars: 30 μm.

## Discussion

The rapid tip growth and rupture of the pollen tube are believed to result from a sophisticated control mechanism between the elastic properties of the pollen tube tip cell wall and turgor pressure (Dehors et al., 2019; Cascallares et al., 2020; Hill et al., 2012; Weigand et al., 2023; Winship et al., 2010). This study showed that pollen tube rupture can be induced by blue light irradiation. This novel photo-regulatory burst can serve as a useful technique to monitor the rapid alteration of sperm cells, which typically occurs immediately after pollen tube discharge.

### How does blue light cause Ca^2+^ influx?

In this study, we observed enhanced Ca^2+^ influx in most pollen tubes after blue light irradiation (Figure 3). Based on the fluorescent pattern of GCaMP and photosensitive region (Figure 3, 4), Ca^2+^ influx appears to be caused by calcium channels localized on the pollen tube apex. The *Arabidopsis* genome encodes at least 77 candidate calcium-permeable channels, including CNGCs, GLRs, TPCs, MSLs, OSCAs, a piezo-type channel, and annexins. Among them, at least 39 channels were expressed in pollen (Johnson et al., 2019), and a subset of the calcium channels (i.e., CNGC18, MLO1/5/9/15) were localized to the pollen tube tip (Chang et al., 2007; Gao et al., 2016; Gao et al., 2023; Meng et al., 2020). Some of the tip-localized calcium channels are controlled by ligands, such as cAMP, cGMP, and RALF peptides. Blue light irradiation may induce environmental changes that alter the associations of these channels and ligands. Alternatively, blue light irradiation can induce abiotic signals like ROS. ROS can induce Ca^2+^ influx and is indispensable for Ca^2+^ gradient formation at pollen tube tips (Duan et al., 2014).

In addition to calcium channels and their agonistic signals, the photoreceptor poses the biggest mystery in blue light-induced Ca^2+^ influx. Several photo-activated proteins, including PHOTOTROPINs, CRYPTOCROMEs, ZEITLUPE family proteins, and PAS/LOV proteins, respond to the blue light signal (Ogura et al., 2008; Ito et al., 2012). Except for ZEITLUPE (AT5G57360), these proteins exhibit low expression levels in pollen (Winter et al., 2007). However, as pollen tubes grow in deep maternal tissues such as the pistil, they are unlikely to receive light signals *in vivo*. The physiological significance of photosensitivity in pollen tubes remains unknown. Recently, Cho et al. (2023) proposed light-induced partial membrane disruption and tiny pore formation that elevate membrane permeability to cations and a fluorescent dye in mammalian cells. While direct evidence of transient pore formation is still lacking, the model of light-induced membrane permeability change offers a compelling explanation for the immediate enhancement of Ca^2+^ influx. Further analyses focusing on calcium channels, ROS, photoreceptors, or dynamic changes in the phospholipid bilayer at the pollen tube tip may clarify the mechanism of Ca^2+^ influx in response to blue light irradiation.

### Physiological effect of rapid Ca^2+^ influx during pollen tube growth

In our live imaging, inhibition of pollen tube growth or rupture occurred immediately after blue light-induced Ca^2+^ influx (Figure 3). The Ca^2+^ influx at the pollen tube tip is closely linked to the ANX/BUPS/LLG/RALF signaling module and the LRX/RALF module, essential for pollen tube growth and discharge, controlling cell wall integrity (Gao et al., 2023; Fabrice et al., 2018; Ogawa and Kessler, 2023). For instance, the absence of Ca^2+^ influx through the Calcium channel MLO1/5/9/15, triggered by RALF4/19 peptides binding to the ANX/BUPS/LLG complex, disrupts cell wall integrity, leading to pollen tube burst (Gao et al., 2023). Earlier studies indicated that Ca^2+^ ionophore treatment increases pollen tube cell wall thickness (Reiss and Herth, 1979; Picton and Steer, 1983). Considering these findings, pollen tube burst in the absence of Ca^2+^ influx in the mutant of the ANX/BUPS/LLG complex may result from decreased pollen tube cell wall thickness, explaining the rupture as an imbalance between cell wall strength and turgor pressure.

Conversely, we observed pollen tube burst with an increasing cytosolic Ca^2+^ level after blue light irradiation (Figure 3). The prerupture Ca^2+^ influx was also noted in semi-*in vivo* fertilization conditions (Ngo et al., 2014). Additionally, a similar Ca^2+^ pattern was reported in ROS-inducible burst and spontaneous pollen tube burst in the *lrx8/9/11* triple mutant in *in vitro* conditions (Duan et al., 2014; Fabrice et al., 2018). Most likely, excess Ca^2+^ influx could also induce an imbalance between cell wall stiffness and turgor pressure, but the precise downstream effects are still debated. Calcium channel opening may cause a sudden water influx, increasing turgor pressure (Feher and Lajko, 2015). Alternatively, a higher Ca^2+^ level could accumulate ROS at the pollen tube tip via the activation of RESPIRATORY BURST OXIDASE HOMOLOG H (RBOHH) and RBOHJ (Kaya et al., 2015), potentially weakening the pollen tube cell wall and promoting rupture (Feher and Lajko, 2015).

Interestingly, 66% of pollen tubes did not rupture, despite strong Ca^2+^ influx induced by blue light irradiation (Figure 3 F). The absence of pollen tube burst may simply result from Ca^2+^ influx levels below the threshold of rupture execution. However, considering these nonruptured pollen tubes stopped tip growth, a subset of pollen tubes under growth arrest might tolerate catastrophic pollen tube burst. In our results, weak blue light irradiation tended to induce growth arrest rather than rupture (Supplemental Movie 2). The apex of these quiescent pollen tubes would halt cell wall synthesis and acquire sufficient stiffness against turgor pressure. Conversely, pollen tubes accelerate tip growth prior to pollen tube discharge in receptive synergid cell (Ngo et al., 2014). Pollen tubes can facilitate their burst by actively generating nascent cell wall. To our knowledge, pollen tube growth and pollen tube burst were independently discussed in physiological and pharmacological studies. Our results elucidate the correlative aspect of growth and burst in the pollen tube and suggest the importance of pleiotropic analyses, including both responses.

### Application of blue light-induced pollen tube rupture

Blue light-induced pollen tube rupture can be a potent tool for analyzing the rapid fertilization mechanism in flowering plants. Double fertilization in *Arabidopsis* is completed within 10 min after pollen tube burst (Hamamura et al., 2011). Subsequent IVPM breakdown and GCS1/HAP2 relocation must occur swiftly before gamete fusion (Sugi et al., 2023; Sprunck et al., 2012; Wang et al., 2022). To observe these events under a microscope, the development of a new technique for controlling pollen tube discharge timing and position was eagerly anticipated. Several methods exist for isolating sperm cells from pollen tubes, such as physically cutting pollen tubes with a surgical blade and treatment with different concentrations of mannitol (osmotic shock), ROS, and RALF34 (Sprunck et al., 2012; Shi et al., 2022; Duan et al., 2014; Ge et al., 2017). However, these often result in sample drifting during dissection or chemical treatments, hindering continuous imaging. In contrast, our blue light-induced method allows for the preparation of fresh sperm cells simply by changing the microscope channel, requires no expensive equipment, and minimizes sample drifting. A continuous imaging of identical sperm cells using our blue light-induced method will elucidate the dynamics of IVPM fragmentation and subtle quantitative changes in GCS1/HAP2 relocation upon pollen tube discharge. These data will provide fundamental knowledge on sperm cell activation and contribute to understanding the double fertilization mechanism at the molecular level.

## Materials and Methods

### Plasmid construction

A binary plasmid containing *LAT52pro:GCaMP* was generated as follows. The *LAT52* promoter sequence was isolated from *LAT52p::Venus* (Mizuta et al., 2015) through *Hin*dIII/*Bam*HI digestion. The *GCaMP6f* sequence was PCR-amplified from *pGP-CMV-GCaMP6f* (#40755, addgene) and cloned into the pCR-Blunt II-TOPO vector (Thermo Fisher Scientific, USA). The *GCaMP6f* fragment was isolated via *Bam*HI/*Xba*I digestion. Both fragments were then cloned into the pPZP211 vector (Hajdukiewicz et al., 1994), containing the Nos-terminator sequence, utilizing *Hin*dIII and *Xba*I sites.

### Plant material and Growth condition

*Arabidopsis thaliana Columbia-0* served as the WT plant. Both WT and *pLAT52:GCaMP* marker seeds were germinated on Murashige-Skoog medium and subsequently transferred to soil. The plants were cultivated at 22°C under continuous lighting conditions. *N. benthamiana* seeds were germinated on Murashige-Skoog medium, then transferred to soil, and grown at 25°C under long-day conditions (16 h light/8 h dark). *T. fournieri* cv. “Blue and White” was cultivated at 22 °C under long-day conditions (16 h light/8 h dark).

### Time-lapse imaging of pollen tube growing in semi-in vivo condition

In *A. thaliana*, the following medium and experimental technique were employed. Emasculated WT pistils were cut beneath the style using a 27-gauge needle. The pistils were placed on pollen tube growth media (0.01% (w/v) boric acid, 5 mM CaCl_2_, 5 mM KCl, 1 mM MgSO_4_, 10% (w/v) sucrose, 10 mM epibrassinolide, 1.5% (w/v) NuSieve GTG agarose, pH 7.5) cast in a glass bottom dish (D11130H; Matsunami Glass, Osaka, Japan). Subsequently, the stigma was pollinated with WT pollen. After 3 h of incubation at 22°C, pollen tubes emerging from the cut style were used for microscopic observation. For *N. benthamiana*, the medium and experimental technique were the same as in *Arabidopsis* with slight modifications. In the incubation step, approximately 24 h at 25°C were needed for sufficient pollen tube growth. In *T. fournieri*, 12 mm-long hand-pollinated styles were placed on modified Nitsch’s medium (Higashiyama et al., 1998) containing 1.5% (w/v) NuSieve GTG agarose. After 8 h of cultivation in the dark at 22 °C, pollen tubes emerged from the cut pistil.

### Light irradiation during bright field observation

Pollen tubes from *A. thaliana, N. benthamiana*, and *T. fournieri*, growing in a semi-*in vivo* condition, were observed using an inverted microscope (IX-73; Evident, Tokyo, Japan) equipped with a spinning disk confocal scanning unit (CSU-W1; Yokogawa, Tokyo, Japan) and an sCMOS camera (Zyla 4.2; Andor, Belfast, Northern Ireland). Weak transmitted light from a halogen lamp was utilized for illumination, with excitation light from a CFP filter set (U-FCFP; Evident, Tokyo, Japan), GFP filter set (U-FGFP; Evident, Tokyo, Japan), and RFP filter set (U-FGWA; Evident, Tokyo, Japan)(data sheet for the CFP filter set: EX, 425–445 nm, EM 460–510 nm, the GFP filter set: EX, 460-480 nm, EM 495-540 nm, RFP filter set: EX, 530-550 nm, EM 575-625 nm) was used for light irradiation.

### Calcium imaging

The sample preparation is the same as above, except for using *pLAT52:GCaMP* marker pollen. To observe GCaMP behavior when irradiated by excitation light from a CFP filter set, a 488-nm laser (OBIS 488, Coherent, Santa Clara, California) was employed for GCaMP excitation, and a fluorescence light source (U-HGLGPS, Evident, Tokyo, Japan) served as the stimulus for pollen tube burst. The emission filter was removed from the original CFP filter set to allow the fluorescence of GCaMP to pass through. Time-lapse images of the pollen tubes were captured every second.

## Data Availability Statement

Supplemental information includes experimental procedures with movie captions, five videos and can be found with this article online at https://doi.org/xxxxxxx.

## Acknolwedgements

We thank H. Ikeda, and H. Kakizaki for their technical support.

## Author Cntributions

NS and DM designed the study and wrote the manuscript. NS discovered the pollen tube rupture induced by blue right irradiation. YM generated the *pLAT52:GCaMP* marker. DS contributed analysis in *T. fournieri*. NS collected other data used in this manuscript. TK provided critical advice and reviewed the manuscript. All authors have contributed to the manuscript and approved the manuscript submission.

