## Supplementary text for "Blue light irradiation induces pollen tube rupture in various flowering plants"

**Supplementary Movie Legend**

Supplementary Movie 1. Irradiation of CFP filter excitation light in 100% intensity to pollen tubes. Time-laps movie of blue light irradiation to wild-type pollen tubes growing in semi-in vivo condition at 1-s intervals. The irradiation light was continuously irradiated at 100% intensity. Time stamp (mm:ss): the first frame of the blue right irradiation in the middle of bright field observation was designated as 0 s.

Supplementary Movie 2. Irradiation of CFP filter excitation light in 25% intensity to pollen tubes. Time-laps movie of blue light irradiation to wild-type pollen tubes growing in semi-in vivo condition at 1-s intervals. The irradiation light was continuously irradiated at 25% intensity. Time stamp (mm:ss): the first frame of the blue right irradiation in the middle of bright field observation was designated as 0 s.

Supplementary Movie 3. Irradiation of CFP filter excitation light to pollen tubes.

Time-laps movie of CFP filter excitation light irradiation to wild-type pollen tubes growing in semi-in vivo condition at 1-s intervals. The irradiation light at 100% intensity was continuously irradiated for 15 s. Time stamp (mm:ss): the first frame of the blue right irradiation in the middle of bright field observation was designated as 0 s.

Supplementary Movie 4. Irradiation of GFP filter excitation light to pollen tubes.

Time-laps movie of GFP filter excitation light irradiation to wild-type pollen tubes growing in semi-in vivo condition at 1-s intervals. The irradiation light was continuously irradiated at 100% intensity for 15 s. Time stamp (mm:ss): the first frame of the blue right irradiation in the middle of bright field observation was designated as 0 s.

Supplementary Movie 5. Irradiation of RFP filter excitation light to pollen tubes.

Time-laps movie of RFP filter excitation light irradiation to wild-type pollen tubes growing in semi-in vivo condition at 1-s intervals. The irradiation light at 100% intensity was irradiated for 15 s. Time stamp (mm:ss): the first frame of the blue right irradiation in the middle of bright field observation was designated as 0 s.
